# Non-Invasive dsRNA Delivery via Feeding for Effective Gene Silencing in Teleost Fish: A Novel Approach in the Study of Gene Function Analysis

**DOI:** 10.1101/2024.02.19.580956

**Authors:** Xiangyu Gao, Ruiyan Yang, Weihao Song, Yiyang Shen, Hao Sun, Tianci Nie, Xinlu Yue, Zongcheng Song, Jie Qi, Quanqi Zhang, Yan He

## Abstract

RNA interference is a powerful technique for gene silencing, involving post-transcriptional regulation of gene expression. Successful application of RNAi has been demonstrated in various organisms like nematodes, insects, and oysters by ingesting dsRNA-expressing bacteria. In this study, we attempted a non-invasive feeding method in the marine economy teleost fish, *Sebastes schlegelii*. To validate the feasibility of this approach in *S. schlegelii*, we utilized rotifers (*Brachionus plicatilis*), *Artemia nauplii*, and commercial pellet feed as vectors to deliver *Escherichia coli* strain HT115 engineered to express double-stranded RNA (dsRNA), targeting *dead end* (*dnd*) gene, known to be important for primordial germ cells (PGCs) migration and proliferation in fish. Our experimental results consistently showed that knockdown of the *dnd* gene effectively inhibited the development of PGCs in *S. schlegelii*, resulting in gonadal dysplasia and even sex reversal. This study marks a significant advancement in understanding gene function in teleost fish, laying the groundwork for future large-scale research in this field.

## Introduction

RNA interference (RNAi) is a groundbreaking technique that regulates gene expression by inducing gene silencing through dsRNA (Mello and Conte, 2004; Tuschl, 2001). Its discovery in *Caenorhabditis elegans* demonstrated the ability to silence target genes upon the introduction of dsRNA with the same sequences (Fire et al., 1998). With the advent of this technology, its application has also been widely carried out in many other species such as insects (Cooper et al., 2019; Gu and Knipple, 2013; Kong et al., 2014), plants (Bharathi et al., 2023; Zotti et al., 2018), and mammals (Carmell et al., 2003; Song et al., 2003). By regulating the expression of relevant genes, it is of great significance in the study of gene function, yield improvement, and disease prevention. The interference of dsRNA can be accomplished through different methods, such as dipping, applying, injecting, and feeding (Itsathitphaisarn et al., 2017). Although the injection method has been employed successfully in *C. elegans*, *Drosophila*, *Plutella xylostella,* and various mammals, it has notable technical limitations. Both microsyringes and glass syringes unavoidably contact the species’ bodies, increasing the risk of internal organ injuries and potential mortality (Payton et al., 2017). Besides, the technique requires a relatively high level of experimental skill and effort. Consequently, the number of injected individuals is typically limited, hindering large-scale experiments. Fortunately, the non-invasive feeding method emerges as an ideal solution to these challenges. Initially reported in *C. elegans* (Timmons and Fire, 1998), this method has been successfully implemented in insects (Huvenne and Smagghe, 2010; Nunes and Simoes, 2009), oysters (Feng et al., 2019; Payton et al., 2017), and many other species. However, no reported success has been achieved in teleost fish so far.

In this study, we aim to bridge this gap by introducing the non-invasive feeding method to teleost fish. Our research focuses on its potential application in economically important teleost fish, *Sebastes schlegelii*, distributed in the Pacific Northwest. The initial study of RNAi in teleost fish was carried out in the model organism zebrafish. Researchers successfully knocked down growth-related genes, such as the *floating head* (*flh*) and *no tail* (*ntl*), by injecting dsRNA into zebrafish embryos, yielding phenotypes consistent with the expected results (Wargelius et al., 1999). Meanwhile, injecting dsRNA complementary to the *myostatin* gene into zebrafish embryos at the 1-2 cell stage produced morphologically hyperplastic and hypertrophic zebrafish, with the disruptive effect dependent on the injected dsRNA dosage (Acosta et al., 2005). However, the conventional injection method’s limitations hinder its large-scale application. Although theoretically, we can directly feed bacteria to *S*. *schlegelii* larvae, practical considerations arise as larvae primarily consume rotifers and *Artemia nauplii* at 5-40 days post parturition (dpp) and transition to commercial pellet feed after 40 dpp. To overcome this challenge, we formulated a plan to evenly mix bacteria with rotifers and *A. nauplii* feeding solutions during 5-40 dpp and incorporated the bacteria into commercial pellet feed after 40 dpp, ensuring effective ingestion of the target bacteria by the larvae.

To validate this technique, we have chosen to knock down genes related to PGCs migration. It has been shown that the PGCs of *S. schlegelii* migrate to the germinal ridge at about 10 dpp to form the primordial gonads and differentiate into oogonia at 30 dpp and spermatogonia at 70 dpp (Zhou et al., 2020). As a specific germplasm component first identified in zebrafish, *dnd* has been proven to be crucial to the formation and migration of PGCs in various teleost fish (Li et al., 2016; Weidinger et al., 2003; Yoshizaki et al., 2016). The absence of *dnd* can lead to gonadal dysplasia. Morpholino oligonucleotides (MO) knockout of *dnd* in *Atlantic cod* leads to the loss of PGCs and serves as a sterile receptor for the transplantation of donor spermatogonia (Skugor et al., 2014). Consistently, electroporation of *dnd* knockout constructs into *carp* embryos results in transgenic sterility (Su et al., 2014).

In this work, we utilized the non-invasive dsRNA delivery via feeding technique to knock down the *dnd* gene during PGCs migration and differentiation in the viviparous teleost fish, *S. schlegelii*. Remarkably, the knockdown of the *dnd* gene was proved highly effective in inhibiting PGC development, leading to gonadal dysplasia and, in some cases, even sex reversal. This groundbreaking success in establishing RNAi technology by feeding in teleost fish opens up new avenues for extensive exploration of the gene functions that hold promising potential in fish disease prevention.

## Materials and Methods

### Ethics Statement

This study was approved by the College of Marine Life Sciences, Ocean University of China Institutional Animal Care and Use Committee on Dec 1, 2021.

### Fish

All fish used in this study were obtained from Shenghang Ocean Science and Technology Co., Ltd. (Weihai, Shandong Province). Larvae were reared in circular water tanks measuring 1.5m by 1m, each containing approximately 6000 larvae. We set up a total of three groups, respectively named knockdown group, control group, and blank group. During the early feeding stage, larvae were fed with hatched live rotifers from 5 dpp. As they grew, the larvae were fed with *A. nauplii* at around 20 dpp and commercial pellet feed at about 40 dpp. The seawater temperature, influenced by incoming seawater and sunlight intensity, was maintained at approximately 18 ± 1.5°C. Considering the critical period of PGC differentiation from birth to 70 dpp, we set the feeding cycle at 5-95 dpp. Samples were taken every 10 days, and weight and length were measured simultaneously. To facilitate sampling due to the small size of gonads, we opted to cut off the head and tail and fix the trunk in Bouin’s solution for sampling. After the feeding cycle, fish from different developmental stages were tracked and sampled to confirm the success of the knockdown experiment in various ways. We selected two periods, 116 dpp and 167 dpp, when the gonads could be individually isolated. Half of the gonads were snap-frozen in liquid nitrogen and stored at -80°C for RNA extraction, and the other half were fixed in 4% paraformaldehyde (PFA) for immunofluorescence analysis.

### Vector construction and dsRNA preparation

The L4440 plasmid has a T7 promoter at each end of the multiple cloning site (MCS), which is transformed into the HT115 competent strain and can express dsRNA after IPTG induction. Due to the lack of RNase III in HT115, it is unable to digest RNA. The intact dsRNA is fed into the *S. schlegelii.* Then, the Dicer endonuclease of *S. schlegelii* digests dsRNA to form siRNA, which binds to the target mRNA to perform gene silencing.

The siRNA location was predicted based on the open reading frame (ORF) of *dnd* using the siRNA prediction website http://sidirect2.rnai.jp/ (Sarker et al., 2017). To avoid off-target effects, the IDT PrimerQuest Tool was used for PCR primers design in the region with numerous interference sites (Wang et al., 2020), determining the final size of the interference fragment to be 423 bp. *S. schlegelii* ovarian cDNA was used as a template and enzyme cleavage sites (XbaI and HindIII) were incorporated into the forward and reverse primers to amplify the target fragment. Subsequently, the target fragment was inserted into the PMD-19T vector and transformed into *E. coli* DH5α. The bacterial solution was spread on LB plates (containing 100 mg/L ampicillin and 12.5 mg/L tetracycline) and incubated at 37L for 14 hours. Single colony sequences were obtained by Sanger sequencing. Then, the target fragment was inserted into the L4440 vector and transformed into *E. coli* HT115 (AngYubio, Shanghai, China) competent strain. Then, the single colony was cultivated in a 50ml LB liquid medium containing 100 mg/L ampicillin and 12.5 mg/L tetracycline, and expanded until an OD value between 0.4-0.6 was reached. IPTG (Solarbio, Beijing, China) was added to a final concentration of 0.5 mM and the expression was induced at 37L for 4 h. This produced the treatment group bacterial solution. The HT115 strain, transformed with L4440 empty plasmid and induced by IPTG, was used as the control group.

### Feeding experiment of dsRNA

The feeding experiment was divided into three groups, in which the blank group was fed with normal food sources, and the knockdown and control groups were fed with the *dnd*-L4440-Ht115 recombinant plasmid strains and the L4440-Ht115 empty plasmid strains, respectively. Larvae feed on rotifers from 5-20 dpp, *A. nauplii* from 20-40 dpp, and commercial pellet feed after 40 dpp. Therefore, we developed an experimental plan in which the induced bacterial solution was centrifuged, resuspended, and mixed into the food at a rate of 1 mL/200 larvae/day to feed the larvae. Specifically, during feeding rotifers or *A. nauplii* suspensions, we mixed the bacterial solution with it before feeding. During the feeding of commercial pellet feed, the resuspended bacterial solution was stirred into the feed, dried naturally, and fed to the larvae.

### Histological examination and staining

The gonads were excised and fixed in Bouin’s solution overnight. Afterward, the residual color was washed off with 70% ethanol. Gonads were serially sectioned at a thickness of 4μm using an RM2016 paraffin microtome and placed onto glass slides. Following overnight exposure at 37°C, the slides were dewaxed in xylene for 20 minutes, rehydrated through a series of ethanol gradients (100%-100%-95%-85%-75%-50%-30%), and washed twice in distilled water. For hematoxylin and eosin staining, tissues were stained with hematoxylin for 15 s and eosin for 4 s. Subsequently, the slides underwent treatment with an ascending series of ethanol (30%-50%-75%-85%-95%-100%), followed by two immersions in xylene. Finally, the slides were sealed with neutral gum. Staining results were observed under a Leica microscope for morphological analysis.

### Immunofluorescence analysis

The gonads were removed, excess fat and abdominal membrane were excised. The tissues were fixed in PFA overnight at 4L and dehydrated in a graded methanol series (30%-50%-70%-80%-90%-95%-100%-100%). Tissues were dehydrated twice in anhydrous ethanol, transparentized with xylene, and embedded in paraffin. Gonads were then serially sectioned at a thickness of 4μm using an RM2016 paraffin microtome and placed onto circular glass slides. The slices were dewaxed with xylene, dehydrated with gradient ethanol, and underwent antigen repair at 95 L. Tissue sections were washed with 1x phosphate-buffered saline (PBS) containing 0.3% Triton-100. After blocking with blocking buffer (10% goat serum, Solarbio) for 1h at room temperature, rabbit anti-v*asa* antibody (1:500, GeneTex, USA) was added and incubated overnight at 4 L. After 12h, the goat anti-rabbit IgG antibody (Solarbio, Beijing, China) coupled with secondary fluorescein isothiocyanate was used for green fluorescence detection. Tissue sections were sealed with a DAPI-containing sealer (Beyotime, Shanghai, China) to visualize the gonadal tissue nuclei and photographed under a fluorescent inverted microscope (Leica, Germany).

### RNA extraction and transcriptome sequencing

Since the gonads of *S. schlegelii* at 116 and 167 dpp are quite small, we extracted RNA from the gonads of both the *dnd* knockdown group and the control group using a trace RNA extraction kit (Genstone biotech, Beijing, China). Twelve new libraries covering two developmental stages were constructed and sequenced for expression analysis post *dnd* knockdown. The RNA-seq data were generated using the NovaSeq 6000 following the 150bp paired-end mode. We used Salmon version 0.7.2 to calculate TPM (transcripts per kilobase million) using default parameters, a more precise way to measure gene expression than RPKM (reads per kilobase million) (Wagner et al., 2012). The final result was visualized using the R package pheatmap to draw a heatmap. Data from the *dnd* knockdown and the control groups were compared by DEseq 2 to identify differentially expressed genes (Love et al., 2014). Differences were considered significant when log_2_Foldchange ≥ 1 and *P* value ≤ 0.05.

### Statistics Analysis

Significant differences in body length between different treatment groups were determined using a one-way ANOVA followed by Least Significant Difference multiple comparisons (*P* < 0.05). These difference analyses were carried out using SPSS 26.0.

## Results

### Construction of *dnd*-L4440 recombinant vector

RNA was extracted from the ovarian tissue of *S. schlegelii* and used to generate a cDNA template for PCR amplification. The resulting product showed the expected *dnd* fragment size of 423 bp (Figure 1b, lane 1). The amplified fragment was successfully ligated to the pMD19T vector and transformed into *E.coli* Trans5α. After colony PCR and sequencing, the *dnd*-pMD19T recombinant vector was obtained. Following double digestion with XbaI and HindIII, the *dnd* fragment was ligated into the L4440 empty vector and transformed into Trans5α. PCR and sequencing confirmed the successful construction of the *dnd*-L4440 recombinant vector. The vector was then transformed into the HT115 competent strain, and induction with IPTG resulted in an additional band of around 400bp in the RNA, the successful induction of IPTG and dsRNA generation of the *dnd* gene by HT115 (Figure 1b, lanes 3-4).

**Figure 1.**
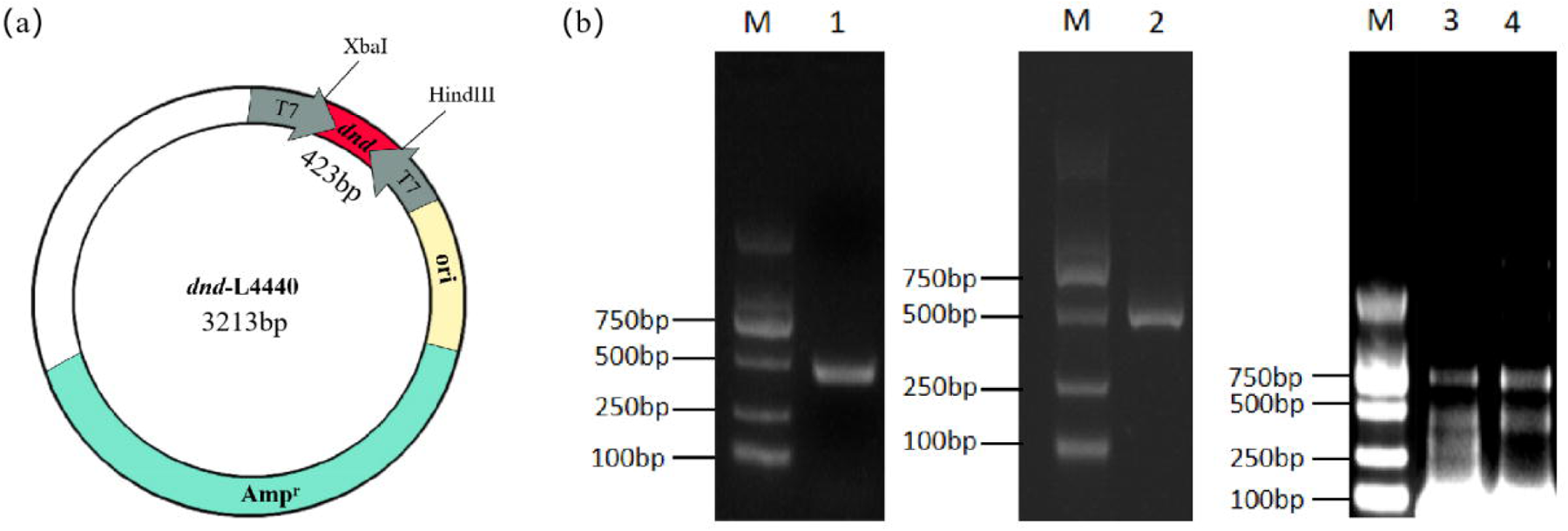
The construction of RNAi plasmid and the induction of dsRNA. (a) Plasmid construction of *dnd*-L4440 with *dnd* fragments sizes and total plasmid sizes. (b) M, DNA marker; 1, PCR products of *dnd* gene fragment; 2, PCR products amplified with *dnd*-L4440 plasmid as a template and M13-F and M13-R as primers. 3, RNA of *dnd*-L4440-Ht115 bacterial solution with IPTG inducer. 4, RNA of *dnd*-L4440-Ht115 bacterial solution without IPTG inducer.

### Growth of *S. schlegelii* was not significantly affected by feeding dsRNA in the long term

During the feeding process, we sampled 45 individuals per group every ten days. In addition, after the feeding experiment, we conducted follow-up sampling at 116dpp and 167dpp respectively. The daily mortality rate could not be accurately counted in any of the three groups during the feeding process because of the large population, but none of them showed large-scale death. We counted the number of surviving individuals of 167 dpp, among which 1630, 1710, and 1650 individuals were in the knockdown group, the control group, and the blank group, with survival rates of 27.2%, 28.5%, and 27.5%, respectively. This suggested that feeding competent strains did not significantly cause the mortality of larvae compared with feeding normal diets. Meanwhile, we regularly measured the weight and body length of each group. Visualization of the results by Prism revealed that during the 90-day feeding cycle, the knockdown group fed with the *dnd*-L4440-Ht115 recombinant plasmid strain showed a significant increase in body length compared to the control group fed with the L4440-Ht115 empty plasmid strain. Specifically, body length was significantly increased (*P* < 0.01) at 40 dpp in the *dnd* knockdown group compared to the control group. Furthermore, this difference reached highly significant at 60 dpp (*P* < 0.0001) and 80 dpp (*P* < 0.0001) (figure 2b). This difference can be clearly distinguished from the appearance (figure 2a). We speculated that the possible reason was that the knockdown of the *dnd* gene inhibited the early migration of PGCs, which in turn affected their differentiation into other germ cells. As a result, some of the energy used for reproductive development was diverted to growth traits, eventually leading to higher body length. It is worth noting that although the survival rate of larvae was not affected by feeding strains, the comparison between the control group and the blank group revealed that growth traits such as body length were significantly reduced when the strains were fed compared to the normal diets, such as at 40dpp (P<0.05) and 60dpp (p<0.0001) (figure 2b). This revealed that feeding bacteria can inhibit the growth of larvae to some extent.

After the feeding experiment, we conducted regular follow-up sampling. The body length of 167 dpp individuals was also measured. The results showed that there was no significant difference in traits between the *dnd* knockdown group and the control group at this time (figure 2b). It is believed that after the feeding experiment, both groups of individuals were fed normal diets, which greatly reduced the interference effect and led to a balance in the growth traits of the two groups. In conclusion, *dnd* knockdown led to an increase in growth traits such as body length in the early stages, while there was no difference in the long term.

**Figure 2.**
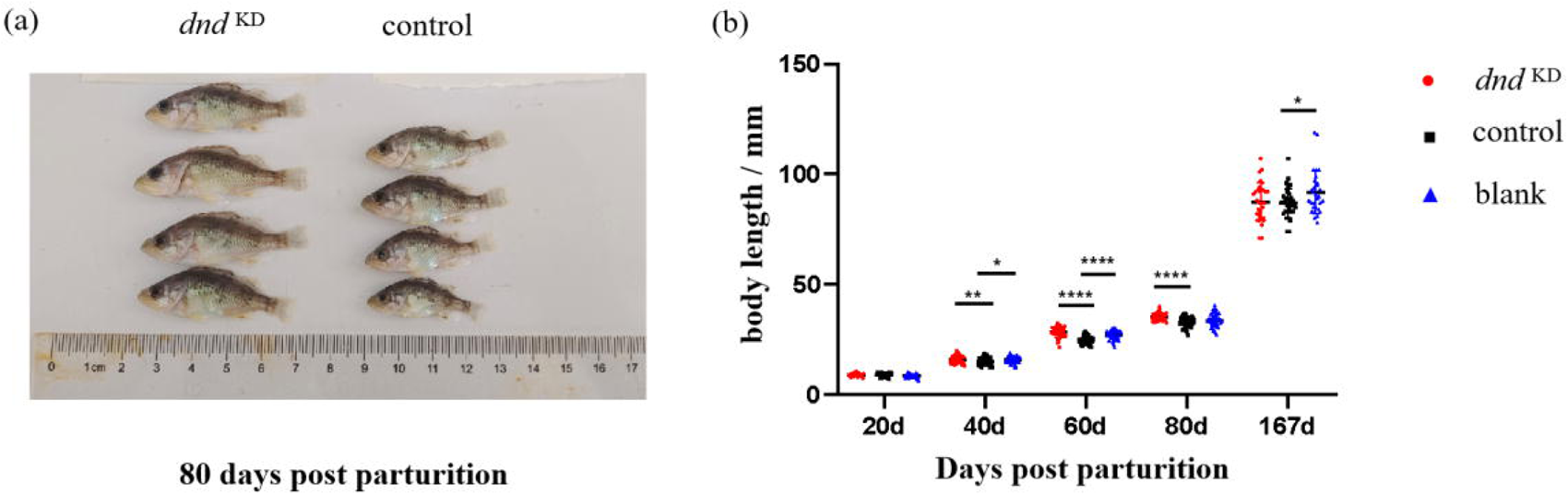
Effects of *dnd* knockdown on body length of larvae at different developmental stages. (a) Comparison of body length between the *dnd* knockdown group and the control group at 80 dpp. (b) Comparison and significance analysis of body length among the *dnd* knockdown group, the control group, and the blank group at different developmental stages (20 dpp, 40 dpp, 60 dpp, 80 dpp, 167 dpp). *, *P* < 0.05; **, *P* < 0.01; ****, *P* < 0.0001.

### *dnd* knockdown altered the expression of genes related to germ stem cells

Compared with the control group, the expression of *dnd* (log_2_FoldChange = -1.075, *P* value = 0.309) in the *dnd* knockdown group of 116 dpp was downregulated (figure 3a). In addition, germ stem cell-related genes such as *vasa* (log_2_FoldChange = -2.034, *P* value = 0.163), *dazl* (log_2_FoldChange = -1.520, *P* value = 0.242), *piwi* (log_2_FoldChange = -1.609, *P* value = 0.173), *sdf1*, *cxcr4*, *nanog*, *gdf3*, *ddx4*, *sox2* were also downregulated (figure 3a), though it was not significant. We further carried out gene expression analysis from the gonads of juvenile fish at 167 dpp. Surprisingly, *dnd* and germ stem cell-related genes were upregulated in the knockdown group (figure 3b). We suspected that there was a negative feedback regulation of *dnd* in the long term after the termination of dsRNA feeding. Interestingly, the knockdown of *dnd* also resulted in sex reversal, where the male-specific gene *amhy* was expressed in physiological females (TPM = 7.05), and the female-specific structural gene *zona pellucida 3* (*ZP3*) was expressed in physiological males (TPM = 567.9).

**Figure 3.**
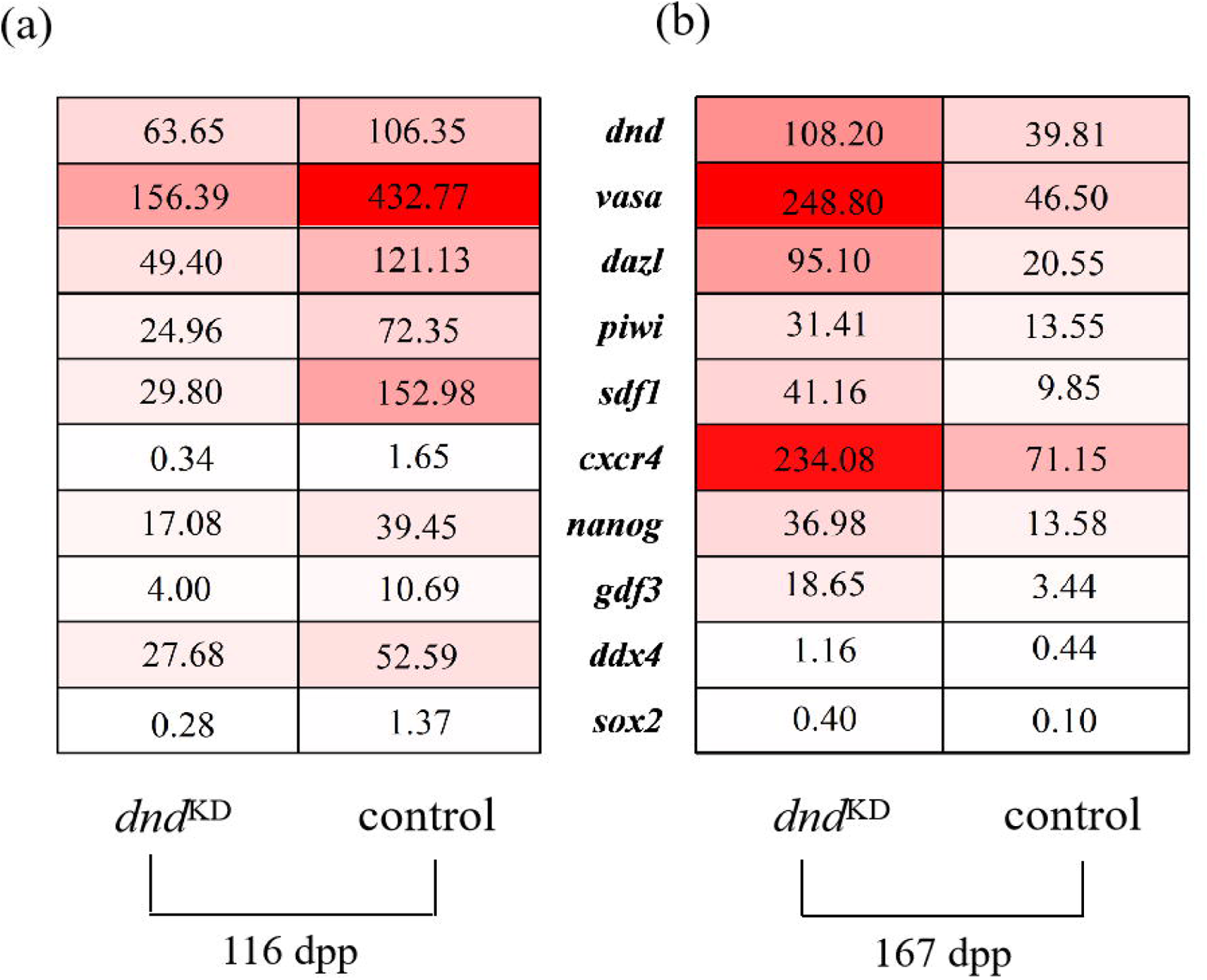
(a-b) Expression patterns of primordial germ cell-related genes at 116 dpp and 167 dpp in two groups of gonads. Numbers indicate the TPM for each gene. Abbreviations: *dnd,* dead end; *vasa, vasa* RNA helicase; *dazl,* deleted in azoospermia-like; *piwi,* P-element induced wimpy testis; *sdf1,* stromal cell-derived factor 1; *cxcr4,* chemokine receptor 4; *nanog, nanog* homeobox; *gdf3,* growth differentiation factor 3; *ddx4,* DEAD-box helicase 4; *sox2,* SRY-box transcription factor 2.

### *dnd* knockdown resulted in gonadal dysplasia in *S. schlegelii*

At 80 dpp, the gonads were too small to be isolated, so the whole trunks were fixed in Bouin’s fluid. However, at 167 dpp, clear differences in gonadal development were observed between the *dnd* knockdown group and the control group. In the knockdown group, the testes showed significant atrophy (figure 4b), while the ovaries exhibited cavitation (figure 4d).

**Figure 4.**
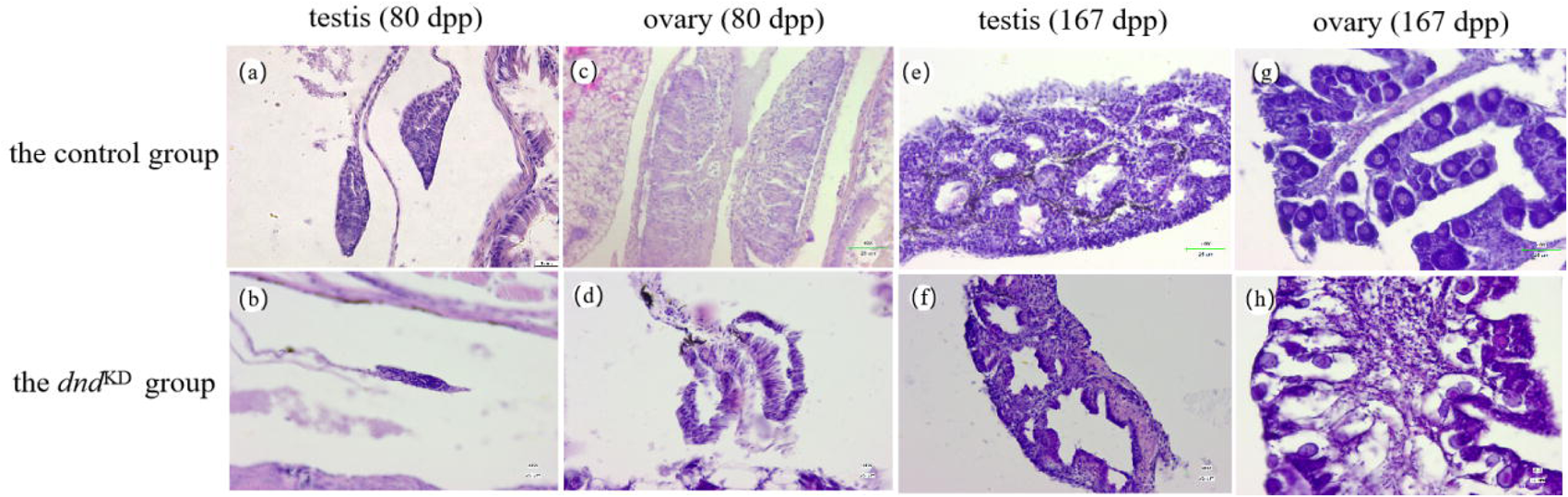
Hematoxylin-eosin staining was performed on the gonads of 80 dpp and 167 dpp. (a) testis of the control group at 80 dpp. (b) testis of *dnd* knockdown group at 80 dpp. (c) ovary of the control group at 80 dpp. (d) ovary of the *dnd* knockdown group at 80 dpp. (e) testis of the control group at 167 dpp. (f) testis of the *dnd* knockdown group at 167 dpp. (c) ovary of the control group at 167 dpp. (d) ovary of the *dnd* knockdown group at 167 dpp. Scale bar = 25 μm.

Out of 60 samples taken from the *dnd* knockdown group, 32 showed gonadal dysplasia. The knockdown group had an ovary that atrophied into a small, sphere-like structure (figure 5h), and the testis showed an uneven distribution of melanin, with one of the pair significantly shorter (figure 5d). In contrast, the gonads of the control group appeared full and uniform in size (figure 5b, f).

Further examination of the internal structures of the gonads revealed differences. In the control group, ovaries contained oogonia and oocytes (figure 4g), while the *dnd* knockdown group showed more fibrous structures in the ovaries (figure 4h). The testes of the control group had numerous spermatogonia, mainly distributed around the outer edge, and spermatic sacs encapsulated by supporting cells were widely present (figure 4e). However, in the *dnd* knockdown group, the testes exhibited a distinct cavity with almost no spermatogonia or seminal vesicles observed (figure 4f).

**Figure 5.**
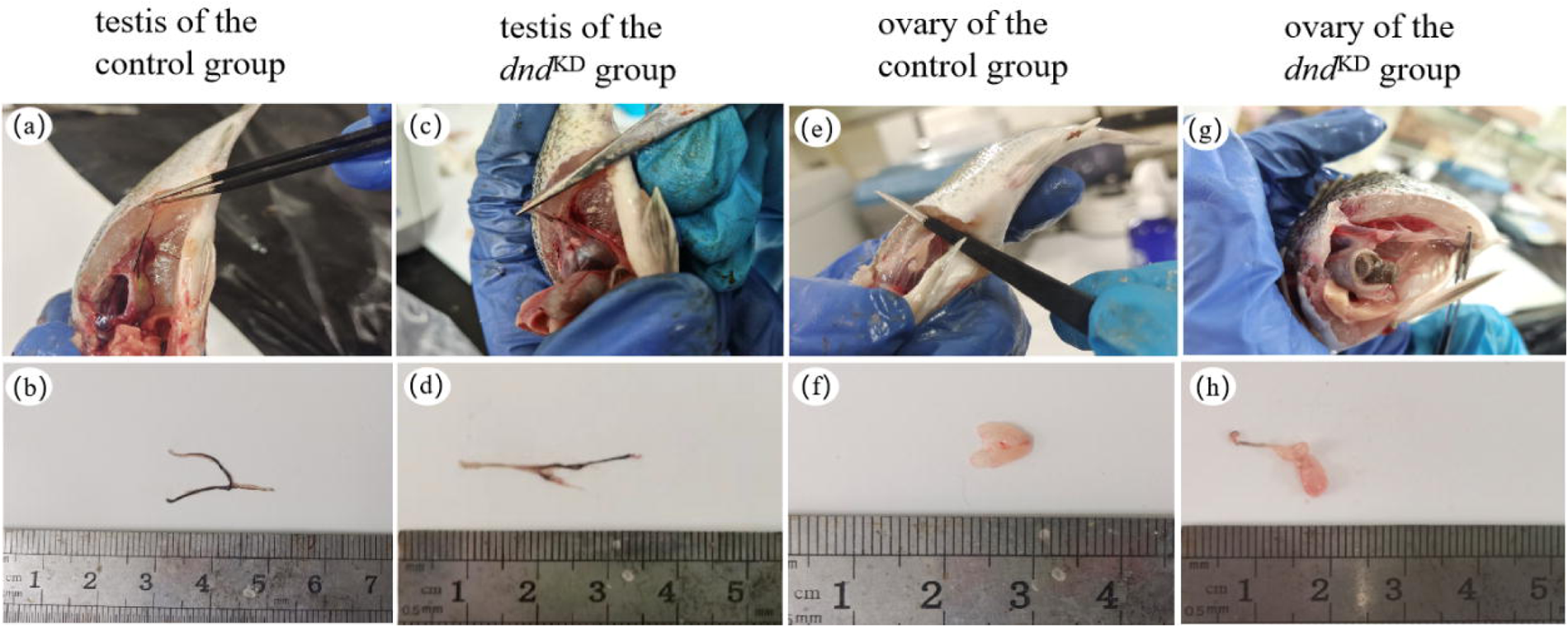
Knockdown of *dnd* leads to dysplasia of ovaries and testes at 167 dpp. (a-b) the appearance of a testis from the control group. (c-d) the appearance of a testis from the *dnd* knockdown group. (e-f) the appearance of an ovary from the control group. (g-h) the appearance of an ovary from the *dnd* knockdown group.

Interestingly, there were two cases in the *dnd* knockdown group where both testes and ovaries were present (figure 6a, e). In the first case, half of the ovary contained oogonia and primary oocytes, while the other half showed a cavity structure. Additionally, although seminal vesicles were present in the testis, the number of spermatogonia was very small, and most of them were somatic cells (figure 6c). The genetic sex of this individual was determined to be male. In another case, the testis of the genetic male contained part of the ovarian structures, including oogonia and primary oocytes (figure 6h). In the control group, no hermaphroditic cases were observed out of 60 samples. After sex identification, the genetic sex of both individuals was male.

**Figure 6.**
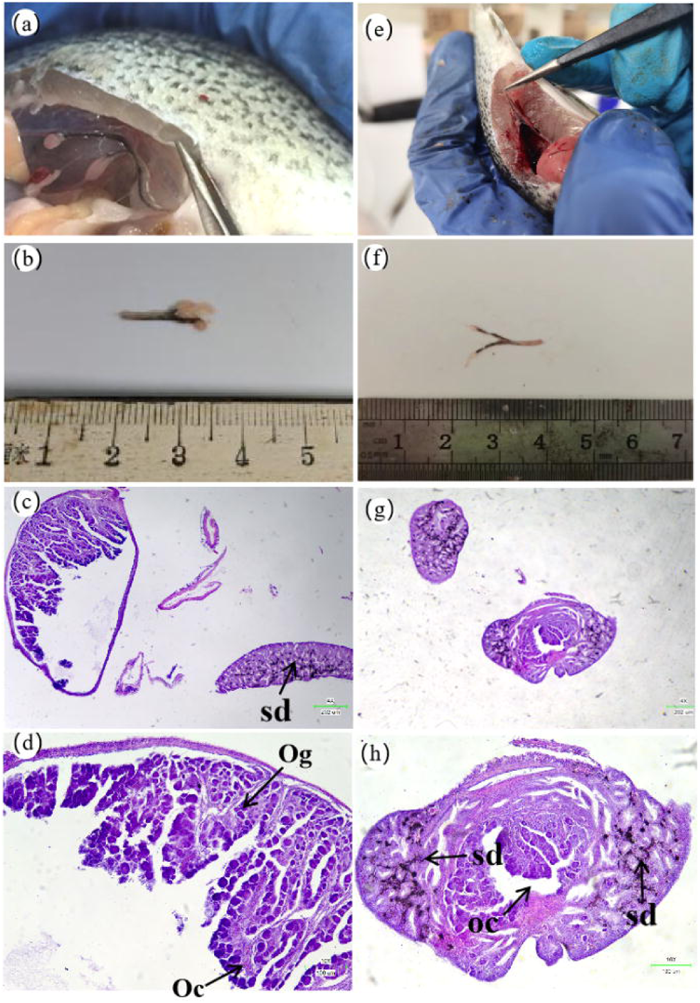
Knockdown of *dnd* leads to hermaphroditic gonads. (a,e) gonads in the *dnd* knockdown group *in vivo*. (b,f) isolated gonads in the *dnd* knockdown group. (c,g) HE staining of hermaphroditic gonads (genetical male). Scale bar = 202 μm. (d,h) HE staining of hermaphroditic gonads (genetical male). Scale bar = 100 μm. Abbreviations: Og, oogonia; Oc, oocyte; oc, ovary cavity; sd, sperm duct.

### *dnd* knockdown resulted in a significant decrease in the number of germ cells

In this investigation, we targeted the marker gene *vasa* in germ cells to explore the protein-level changes after *dnd* knockdown. Immunofluorescence of gonads at 167 dpp revealed that knockdown of *dnd* caused a significant decrease in green fluorescence intensity, whereas the fluorescent signal was strong in the control group. The nuclei in the gonads were stained blue by DAPI to mark cell locations. In the control group, spermatogonia were evenly distributed along the outer margin of the testis (figure 7c). However, in the *dnd* knockout group, the *vasa* protein signal was significantly reduced, and spermatogonia were almost invisible (figure 7d). Similarly, in the ovary of the control group, oogonia and oocytes were evenly distributed (Figure 7g). In contrast, the *dnd* knockdown group exhibited a significant reduction in *vasa* protein signaling, and almost no oocytes were present (Figure 7h).

**Figure 7.**
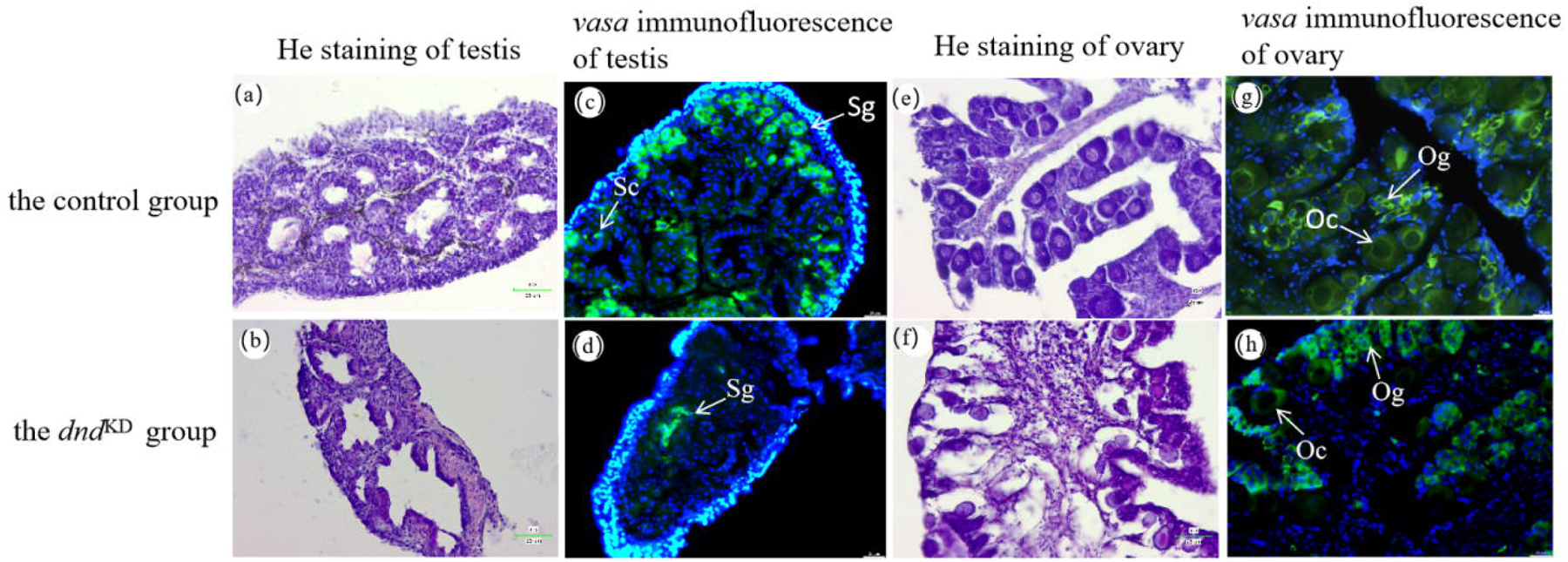
Spatial expression of *vasa* protein in 167 dpp gonads. (a) HE staining of testis in the control group. (b) HE staining of testis in the *dnd* knockdown group. (c) *Vasa* immunofluorescence of testis in the control group. (d) *Vasa* immunofluorescence of testis in the *dnd* knockdown group. (e) HE staining of ovary in the control group. (f) HE staining of ovary in the *dnd* knockdown group. (g) *Vasa* immunofluorescence of ovary in the control group. (h) *Vasa* immunofluorescence of testis in the *dnd* knockdown group. Abbreviations: Sg, spermatogonia; Sc, spermatocytes; Og, oogonia; Oc, oocytes.

## Discussion

This study successfully demonstrated the application of a non-invasive, feed-based RNAi technique in a viviparous teleost fish, *S. schlegelii*. Evidence of this success was presented through the observed gonadal dysplasia in 53.3% of the individuals in the *dnd* knockdown group. In recent years, we have gained an in-depth understanding of fish PGCs migration and proliferation, as well as the timing and mechanism of sex determination. Previous studies on *dnd* in zebrafish (Weidinger et al., 2003), medaka fish (*Oryzias latipes*) (Hong et al., 2016), Atlantic salmon (*Salmo salar*) (Guralp et al., 2020), carp (*Cyprinus carpio*) (Su et al., 2014), and among other teleost fish, has revealed that *dnd* can affect the migration and development of PGCs, leading to gonadal dysplasia. Due to its functional conservation and easily observable phenotypes, we chose the *dnd* gene as a validation gene for establishing RNAi technology. In this study, we performed knockdown experiments for up to 90 days to maximize RNAi effects and phenotypes. Our data suggest that knocking down the *dnd* gene through RNAi with non-invasive feeding methods can obtain the expected phenotype, impaired PGCs formation and gonadal dysplasia. In our previous studies, we have performed RNAi using the same procedure targeting other genes. We have focused on *amhr2*, a type II receptor for the sex-determining gene *amhy* of *S. schlegelii*, and the growth-specific muscle stem cell marker gene *meox1*. After early feeding experiments, we obtained the expected phenotype: individuals with sexual reversal of knocking down *amhr2* and inhibited muscle growth of knocking down *meox1*. These experiments yielded phenotypes consistent with expected gene functions, supporting the specificity and effectiveness of our RNAi method. These two studies have resulted in an approved Chinese patent (No. ZL202211538047.4) and formed the core content of a master’s dissertation (Regulatory role of *meox1* in muscle growth of *Sebastes schlegelii*, master dissertation of Kejia Huang, Ocean University of China). While our study primarily focused on *S. schlegelii*, we posit that the RNAi delivery method demonstrated herein has potential adaptability to other teleost species. The key lies in identifying the critical period when the target gene plays a role in larvae and feeding them a diet mixed with dsRNA, which will enable large-scale gene knockdown in teleost species. This approach not only broadens the applicability of our method but also offers a scalable solution for gene manipulation in aquatic organisms.

Previous studies for gene knockdown in teleost fish utilized dsRNA injection, which was first successfully established in zebrafish. Injecting floating head (*flh*) and no tail (*ntl*) dsRNAs into 1-2 cell stage zebrafish embryos can interfere with the expression of key genes involved in spinal cord development and somite pattern formation, resulting in individuals with developmental abnormalities (Wargelius et al., 1999). Intraperitoneal injection of *igf3*-dsRNA, a gonadal development-related gene, into C*yprinus carpio* was effective in reducing the expression of this gene (Song et al., 2019). In addition, a significant down-regulation of expression was also found when 14-3-3 beta/alpha-dsRNA was intramuscularly injected into *Scophthalmus maximus* (Liu et al., 2020). The injection method, through precise manipulation of a single embryo or individual, typically has high efficiency and can even reach over 80% (Liu et al., 2020; Acosta et al., 2005). In our study, the rough knockdown efficiency is about 50%. Therefore, in terms of knockdown efficiency, bacterial-mediated delivery of dsRNA is not as effective as direct injection (Timmons and Fire, 1998). However, direct contact with the subject through microinjection can cause non-specific defects in zebrafish embryos (e.g. curved tail, gastrulation defects) (Wargelius et al., 1999), and even a higher mortality rate (Song et al., 2019). The non-invasive feeding method we have established in teleost fish can effectively avoid direct contact with the subjects (Lezzerini et al., 2015), greatly reducing its mortality rate and minimizing the adverse effects of interference. Meanwhile, the injection method tends to have fewer subjects because the procedure is more complicated (Song et al., 2019). The feeding method, on the contrary, with its simple operation procedure, can interfere with more experimental subjects, which is conducive to carrying out large-scale experiments.

In the fish surrogacy, it is crucial to produce sterile recipients. Currently, there are three main approaches to generating sterile recipients, including artificial induction of triploid (Piferrer et al., 2009; Takeuchi et al., 2018), interspecies hybridization (Hamaguchi and Sakaizumi, 1992; Shimada and Takeda, 2008), and the use of relevant hormones to hinder gonadal development (Lacerda et al., 2010; Majhi et al., 2014). In *Takifugu rubripes*, fertility can be eliminated through cold shock treatment to induce triploids (Hamasaki et al., 2013). However, the process of triploid induction requires precise timing of the expulsion of the second polar body and optimal survival rate for the zygote, which necessitates high-level technical skill and knowledge. Interspecies hybridization technology is suitable for large-scale production of recipients, but it requires substantial time and economic investment (Ling et al., 2012). In addition, the use of chemical drugs such as busulfan can quickly induce individuals to become sterile (Majhi et al., 2009). However, this method can lead to ulceration and even death in females. Besides, the method is reversible and temporary, unable to maintain infertility for extended periods. This poses a great challenge in controlling the dosage of the drug and choosing the optimal time for interference. These drawbacks largely limit the applicability of these methods in aquaculture.

The PGCs of *S. schlegelii* migrate to the genital ridge to form the primitive gonads at about 10 dpp. They then differentiate into oogonia at 30 dpp and spermatogonia at 70 dpp (Zhou et al., 2020). The *dnd* gene has been proven to be crucial for the survival and migration of PGCs in numerous teleost fish (Li et al., 2016; Weidinger et al., 2003; Yoshizaki et al., 2016). Therefore, we targeted gonadal sterility by knocking down *dnd* at the critical period of PGCs migration and differentiation (5-95 dpp). This method is cost-effective and poses less environmental impact by regulating the gene expression level of the species itself to obtain sterile individuals (Ciruna et al., 2002). Concerning long-term impacts, we have been raising the fish in both the knockdown group and the control group for a year and a half. Our findings indicate that consuming the induced strain at the larvae stage does not significantly affect the survival rates of *S. schlegelii* compared to a normal diet. Up to this point, no adverse effects on fish health and behavior due to bacterial ingestion have been observed. This method, characterized by its major advantages such as scalability, cost-effectiveness, high efficiency, and minimal impact on both organisms and the environment is crucial for gene function analysis in economic fish species. It has the potential to significantly boost the economic viability of fish production by manipulating candidate genes related to economic traits. Furthermore, by increasing the concentration of the induced bacterial solution, we can upscale the RNAi process. This implies that our technology can be effectively integrated into commercial aquaculture, a move of substantial importance. It holds great promise not only for advancing the study of functional genes in teleost fish but also for improving the production of economic fishes. The successful construction of individuals with gonadal dysplasia not only confirms the role of the *dnd* gene in the PGC proliferation and differentiation in *S. schlegelii* but also lays the groundwork for germ stem cell transplantation in this species. Germ cell transplantation (GCT) is a powerful reproductive technology pioneered by Brinster et al. in 1994. By transplanting spermatogonia stem cells from donor mice into the sterile recipient, the recipient will recover fertility and produce functional spermatozoa (Brinster and Zimmermann, 1994). Although GCT is relatively mature in mammals, this technology in teleost fish emerged in 2004, established in salmon (Takeuchi et al., 2004). The GCT of fish is usually carried out during the embryonic period (Ciruna et al., 2002) or the hatching period of larvae (Zhou et al., 2021) when the rejection reaction is relatively small and the success rate of transplantation is high. However, as mentioned earlier, delicate microscopic manipulation is required due to the relatively small receptors during this period, which requires more stringent experimental skills. In addition, sterile recipients constructed by *dnd* knockdown are also susceptible to accepting donor germ cells because the depletion of their germ cells reduces its rejection of exogenous germ cells, facilitating the successful colonization of exogenous germ cells. Transplantation techniques at this time do not require micromanipulation, so the requirement for experimental skills is greatly reduced. This work produces ideal sterile recipients for PGC implantation in *S. schlegelii*.

Another intriguing observation is the male-to-female sex reversal upon the knockdown of the *dnd* gene. Our evidence rests on two main observations. Firstly, upon analyzing gene expression at 116 dpp, we found that the male-specific gene, *amhy*, was expressed in physiological females. Conversely, the female-specific expression gene, *ZP3*, was evident in physiological males. Secondly, at 167 dpp, from a sample of 60 knockdown individuals, we identified two hermaphrodites. It is worth noting that the genetic sex of these examples is male. Therefore, knockdown of the *dnd* gene can lead to a reversal of male to female in *S. schlegelii*, which has not been reported in previous studies on this gene. Interestingly, mutations in the *fancl* gene in zebrafish (Rodriguez-Mari et al., 2010; Rodriguez-Mari and Postlethwait, 2011) lead to a decrease in the number of germ cells. Consequently, the somatic cells in the mutated gonads down-regulate the expression of genes associated with ovarian development. This chain of events culminates in the conversion of females to pseudo-males. Such findings suggest that zebrafish sex is influenced by the number of PGCs. In *S. schlegelii*, the *amhy* gene plays a pivotal role in determining sex (Song et al., 2021). Our study reveals that *dnd* gene knockdown similarly induces sex reversal in this species. This evidence strongly indicates that the number of PGCs influences the sex determination of *S. schlegelii* as well. The interplay between *amhy* and the number of PGCs in establishing the sex of *S. schlegelii* warrants further research.

In addition, to further improve the efficiency of RNAi, we can optimize the feeding experiment. Since the current feeding dose had minimal impact on the survival of larvae, we could increase the bacterial feeding dose to ensure that more dsRNA could be consumed by each individual. Moreover, the average total length of 95 dpp individuals is 53.38 mm, at which stage the individuals were too small to dissect their gonads. We can only dissect the individual gonad as early as 116 dpp (average total length of 72.99 mm) to dissect individual gonads for RNA extraction. Although the expression of *dnd* and PGCs-related genes were still down-regulated at this time, the significance did not reach the expected level since the feeding cycle had been over for 20 days.

In summary, this study has effectively shown that the technique of feeding dsRNA-producing *E. coli* for RNAi is feasible in the teleost fish, *S. schlegelii*. Moving forward, this non-invasive method can be applied to explore the gene function in teleost fish on a large scale. In addition, the hermaphroditism caused by *dnd* knockdown and the successful construction of the sterile recipients offers new directions for the study of fish sex reversal genes and GCT.

## Data accessibility

Data supporting the findings of this study are available within the paper and electronic supplementary material. The RNA-seq data in this project will be submitted to NCBI Sequence Read Archive (SRA) under the project number PRJNA1002463.

## Acknowledgements

This study was financially supported by grants from National Key R & D Program of China (2022YFD2400103), National Natural Science Foundation of China (32070515), Key R & D program of Shandong Province (2022LZGC016) and Technological Small and Medium-sized Enterprises Innovation Enhancement Project of Shandong Province (2023TSGC0638). This work was also supported by High-performance Computing Platform of YZBSTCACC. The funders were not involved in the design of the study and collection, analysis, and interpretation of data and in writing the manuscript.

